# TCMNP: a data processing and visualization database and R package for traditional Chinese medicine network pharmacology

**DOI:** 10.1101/2024.07.13.601094

**Authors:** Jinkun Liu, Jing Feng, Bin Wu, Min Ying

## Abstract

**Background:** Network pharmacology is pivotal for the study of the mechanism of action of traditional Chinese medicine (TCM). However, there are currently some challenges in data processing and visualization for traditional Chinese medicine network pharmacology (TCMNP). We develop the R-environment-based software database and package TCMNP.

**Methods:** TCMNP is designed for the analysis and visualization of compositions in TCM compounds, herbs, ingredients, corresponding targets and pathway enrichment. Data analysis and visualization in R using “dplyr”, “clusterProfiler”, “ggplot2”, and “circlize” packages, as well as functions of our own making.

**Results:** TCMNP is designed for the analysis and visualization of compositions in TCM compounds, herbs, ingredients, corresponding targets and pathway. The TCMNP database contains 571 kinds of TCM herbs, 17,118 ingredients, 10,013 diseases, and 15,956 targets. TCMNP simplifies the data analysis process and realizes a complete set of process operations from Chinese herbal compound composition, component and target automatic screening, enrichment analysis visualization, protein interaction and TF and target gene screening. Its built-in functions of various types provide a tidy interface for the visualization of TCM network pharmacology. TCMNP database is freely available at https://tcmlab.top/tcmr. TCMNP package is freely available at https://github.com/tcmlab/TCMNP.

**Conclusion:** The TCMNP database and package provide a comprehensive overview of the interconnections between components of a TCM compound and its potential effects on disease treatment. TCMNP database is available for free at https://tcmlab.top/tcmr/. TCMNP package is available for free at https://github.com/tcmlab/TCMNP. We anticipate that TCMNP will help to explore the mechanism of TCM to treat diseases.

## Introduction

Traditional Chinese Medicine (TCM) has played an important role in the treatment of diseases and the fight against the Corona Virus Disease 2019 (COVID-19)[1]. However, due to its complex composition and targets, deep research of its mechanisms has not been carried out extensively. Network pharmacology is applied to the analysis of TCM components and an exploration of the mechanism of actions of drugs in treating diseases. This is achieved through a functional enrichment analysis based on known component targets [2]. As of April 30, 2024, there have been approximately 56,000 documents retrieved through the PubMed search engine using “network pharmacology” as a keyword. This suggests the large potential for the application of network pharmacology. However, despite its rapid development, the data processing and visualization aspect of network pharmacology has been lagging, which might impede its usage.

R is a freely available software environment for statistical computing and graphic design that is compatible with various Linux platforms, Windows, and MacOS. Currently, some charts in network pharmacology articles can be made using R packages such as “ggplot2” [3], “circlize” [4] and “clusterProfiler” [5]. Yet, these involve data sorting, parameter adjustment, and code modification among other things. Non-expert users might find them time-consuming and challenging to master. To date, there are no specific R packages for network pharmacology as well as interactive databases, necessitating the development of tools specifically targeted at network pharmacology users.

In this work, we have developed TCMNP, a database and R package dedicated to TCM network pharmacology. This database is primarily designed for analysis and visualization of compositions in Chinese medicine compounds, ingredients, corresponding targets, pathway enrichment, protein-protein interaction (PPI) and transcription factors (TFs).

## Materials and methods

### Data collation

TCMNP employs an open-source drug target database and corroborates it with integrated genomic, proteomic, and metabolic databases to ensure the precision of the generated networks. The data for TCMNP network pharmacology are primarily sourced from deeper integration of data, including TCM Systems Pharmacology Database (TCMSP) [6], DrugBank [7], Encyclopedia of TCM (ETCM) [8] and so on. The data of TFs and their corresponding target genes source from the Transcriptional Regulatory Relationships Unraveled by Sentence-based Text mining (TRRUST) database [9], including two species: human and mouse, which have relatively high reliability after manual annotation. The enrichment analysis was accomplished using the “clusterProfiler” package[5] and “GseaVis” (https://cran.r-project.org/web/packages/GseaVis/) in R.

### Herbal and compound screening related to disease targets

A new function named tcm_prescription was added to the TCMNP database and package. The tcm_prescription function by entering the target genes of the disease, treatment herbs and their prescription can be automatically obtained. Then the herbs and prescription present in a table and visually intuitive bar plot.

The main principle of the tcm_prescription function is to use the mathematical principle of hypergeometric distribution.

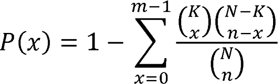

N is the total number of herbs in the background distribution. K is the number of herbs within the distribution that are annotated to the herbal compound of interest. n is the size of the list of herbs of interest and m is the number of herbs in that list annotated to the compound. The default background distribution is all herbs in the database.

### Data visualization

Histograms and dot plots are primarily created using the ggplot2 package in R. The network graph is accomplished utilizing the “igraph” (https://CRAN.R-project.org/package=igraph) and “ggraph” packages (https://CRAN.R-project.org/package=ggraph). The circle graph is predominantly finalized using the “circlelize” package [4]. Sankey and alluvial diagrams are created using the “ggsankey” package (https://github.com/davidsjoberg/ggsankey). Venn diagrams are constructed using the “ggvenn” (https://CRAN.R-project.org/package=ggvenn) and “UpSetR” packages (https://CRAN.R-project.org/package=UpSetR).

## Results

### Main contents of database

TCMNP is divided into nine categories(Figure 1): (1) Content related to traditional Chinese medicine: including herbal medicines, compounds, and targets; (2) Disease information: diseases and disease target genes; (3) Visualization of the composition of traditional Chinese medicine compounds; (4) Kyoto Encyclopedia of Genes and Genomes (KEGG) and Gene Ontology (GO) enrichment analysis; (5) Gene Set Enrichment Analysis (GSEA) enrichment analysis; (6) PPI network interaction system; (7) transcription factor screening and visualization; (8) drug and disease interaction analysis; (9) heat map visualization after molecular docking scoring. The TCMNP database upload file format is a single csv file (Supplementary Table S1). Detailed instructions for using the TCMNP R package and database are provided in Supplementary code S1, Supplementary Videos S1 and S2.

**Figure. 1.**
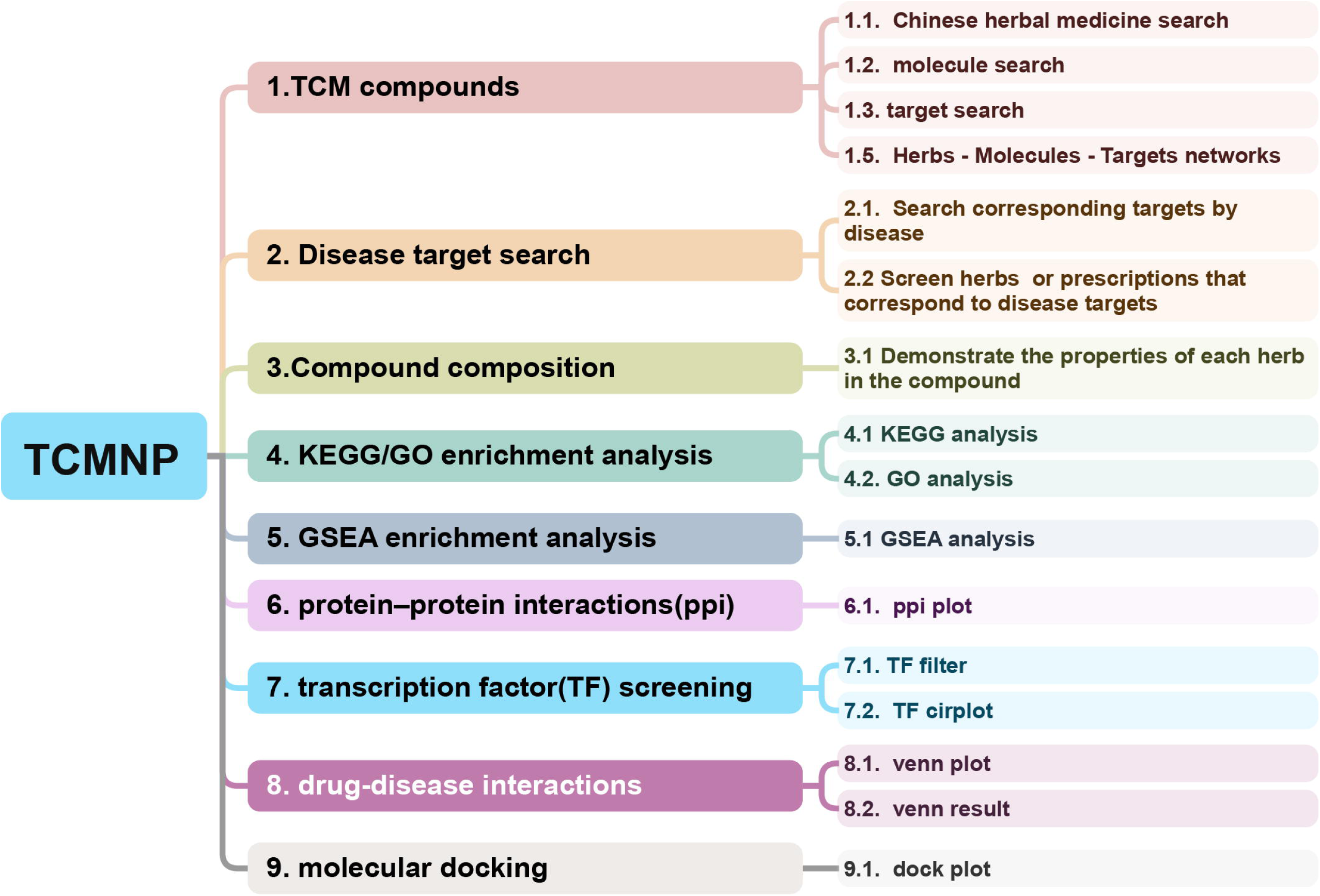
Summary of TCMNP functions.

### Drug target screening

TCMNP database contains 571 kinds of Chinese medicinal materials, 17,118 ingredients, 158 kinds of commonly used prescriptions of TCM and 250 kinds of proprietary Chinese medicines (Figures 2A). TCMNP database can support screening by the names of single or multiple medicinal materials, ingredient names and target names. Network diagrams and Sankey diagrams were produced from the filtered data. In Figure 2B, a Sankey diagram illustrates the herb-ingredient-target interaction. The Sankey diagram of TCMNP database is not limited to the above functions and can support graphical display composed of any number of columns of data. Key nodes in a network diagram often show broader connectivity to their surrounding environment. The Degree parameter allows revealing the number of connections between critical nodes and surrounding nodes (Figure 2C).

**Figure. 2.**
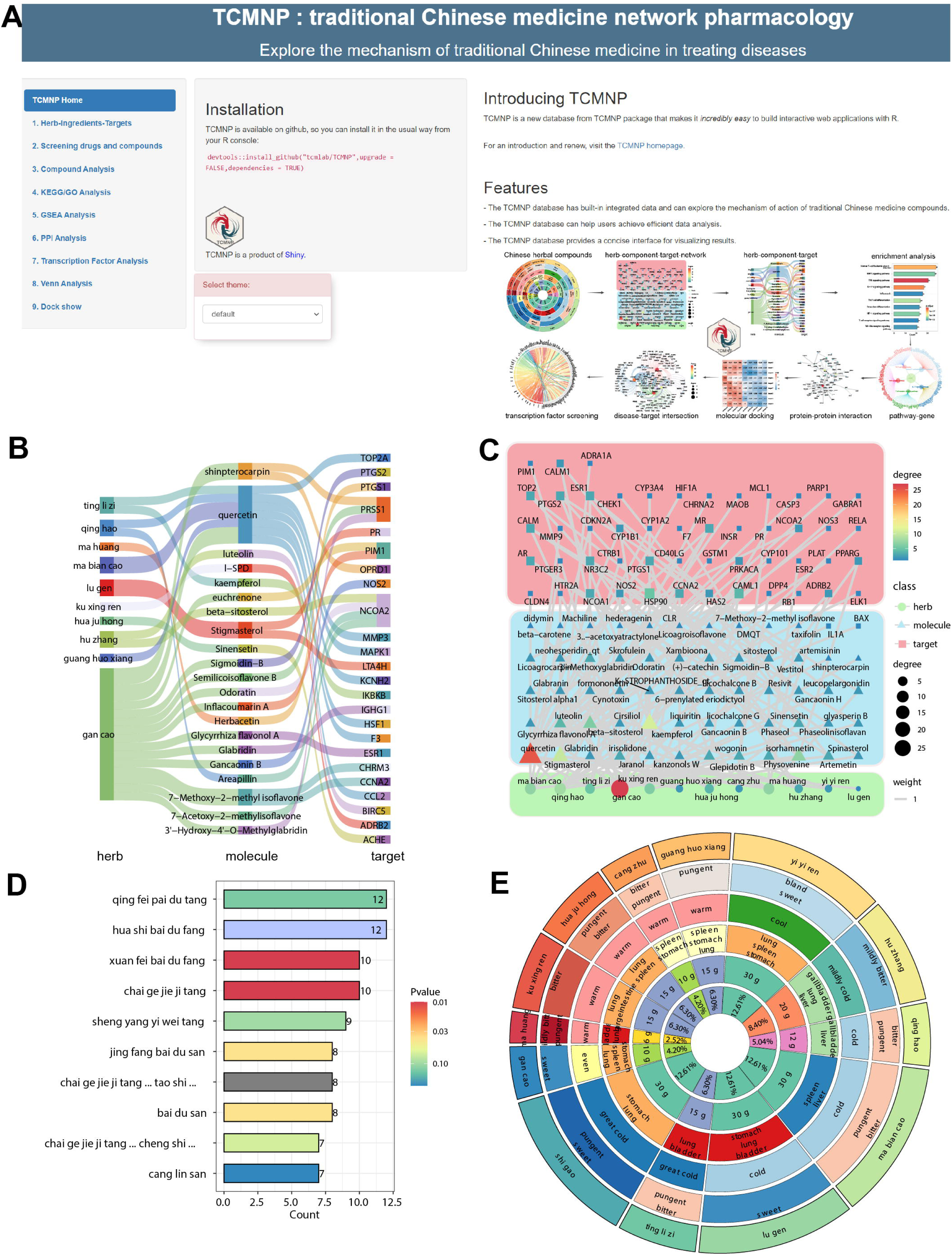
Diagrams related to traditional Chinese medicine compound prescriptions. (A) TCMNP database appearance. (B) Sankey diagrams show the interrelationships between herbs-ingredients-targets. (C) Networks between herbs, herbal ingredients and targets related to them. The degree of a node was defined as the number of its connected edges. The weight expresses the connection times of the relationship. (D) Screening of the top ten prescriptions for treating COVID-19. (E) Xuan fei bai du prescription compound composition. The first circle track indicates the name of the herbs that make up the compound. The second to fifth circles respectively show the herb flavor, herb property, herb meridian tropism and dosage of each herb. The sixth circle and the fan-shaped area indicate the percentage of each herbal medicine dosage.

### Disease target search

We integrated the diseases and targets from the DisGeNET [10] and OMIM [11] databases and obtained a total of 10,013 diseases and 15,956 targets. Users can also summarize disease targets from other data sources and upload them to the TCMNP database, and the background will automatically search for herbal medicines and compound prescriptions related to the disease targets. Take the COVID-19 as an example, the function of tcm_prescription is demonstrated in the Figure 2D. Through analysis, we found that both qing fei pai du decoction, hua shi bai du fang and xuan fei bai du fang (XFBD) were in the top three, which illustrated the accuracy of our new function[1].

### Compound composition analysis

The TCMNP database brings forth a visual representation of the compound composition utilizing the TCMNP package tcm_compound function. These include the herb flavor, herb property and herb meridian tropism of the Chinese medicines. It can be seen from Figure 2E that XFBD main meridians of cold and cool medicines are the lung meridian and bladder meridian, while the main meridians of warm and hot medicines are lung meridian, spleen meridian and stomach meridian. The dose of cold and cool medicine is obviously greater than that of warm medicine. This is consistent with the pharmacological effects of XFBD on antipyretic, antiasthma, anti-inflammation, antiviral and anti-infection [12]. This functional attribute distinctly displays the properties and dosage of various drugs within the Chinese medicinal compounds, thus aiding in understanding the compounds.

### Enrichment analysis

The TCMNP database integrates with the KEGG and GO data by processing with the “clusterProfiler” package. The enrichment results via bar graph (Figures 3A, B), bubble graph (Figure 3C), lollipop graph (Figure 3D) and circle graph (Figure 3E). For exploring pathways, one can employ the pathway circle plot to flaunt essential genes (Figure 3F, Figure 4A). With the help of multiple judgments, we designed all the above functions to be realized through the data format of the data frame and s4 object, which is convenient for later path screening and visualization. As can be seen from Figure 4A, XFBD mitigates COVID-19 symptoms by controlling the expression of pro-inflammatory cytokines like IL-6, IL1β, tumor necrosis factor (TNF), and pathways like IL17A[13]. These articles support such finds.

**Figure. 3.**
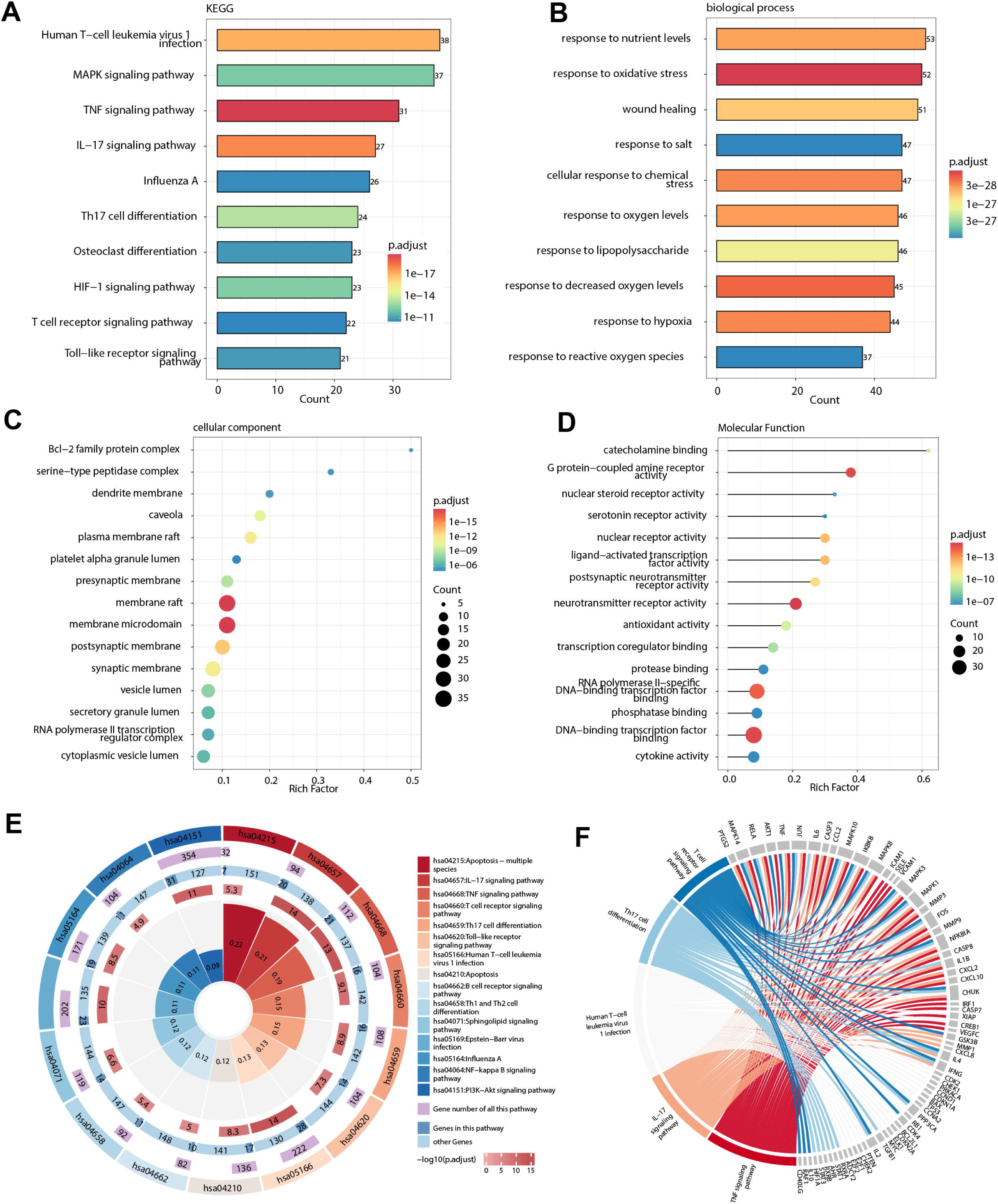
Display of results related to functional enrichment analysis. (A-B) The bar diagrams show the enrichment analysis based on KEGG. (B-D) The GO pathways enrichment analysis results were displayed in the form of bar chart, bubble chart and lollipop chart, respectively. Rich Factor represents the enrichment ratio of genes in the corresponding pathway. The size of the point indicates the genes enriched on this pathway, and the color is presented as -log10 (adjusted *P*-value). (E) KEGG enrichment circle diagram. The first circle represents the pathway id; the second circle represents all the genes in the pathway; the dark color in the third circle represents the enriched genes in the corresponding pathway, and the light color represents the number of other genes involved in this enrichment. The depth of the color in the fourth circle corresponds to -log10 (adjusted *P*-value); the fifth circle (polar coordinate histogram) is the rich factor. (F) Circle diagram showing pathways and the genes on them.

**Figure. 4.**
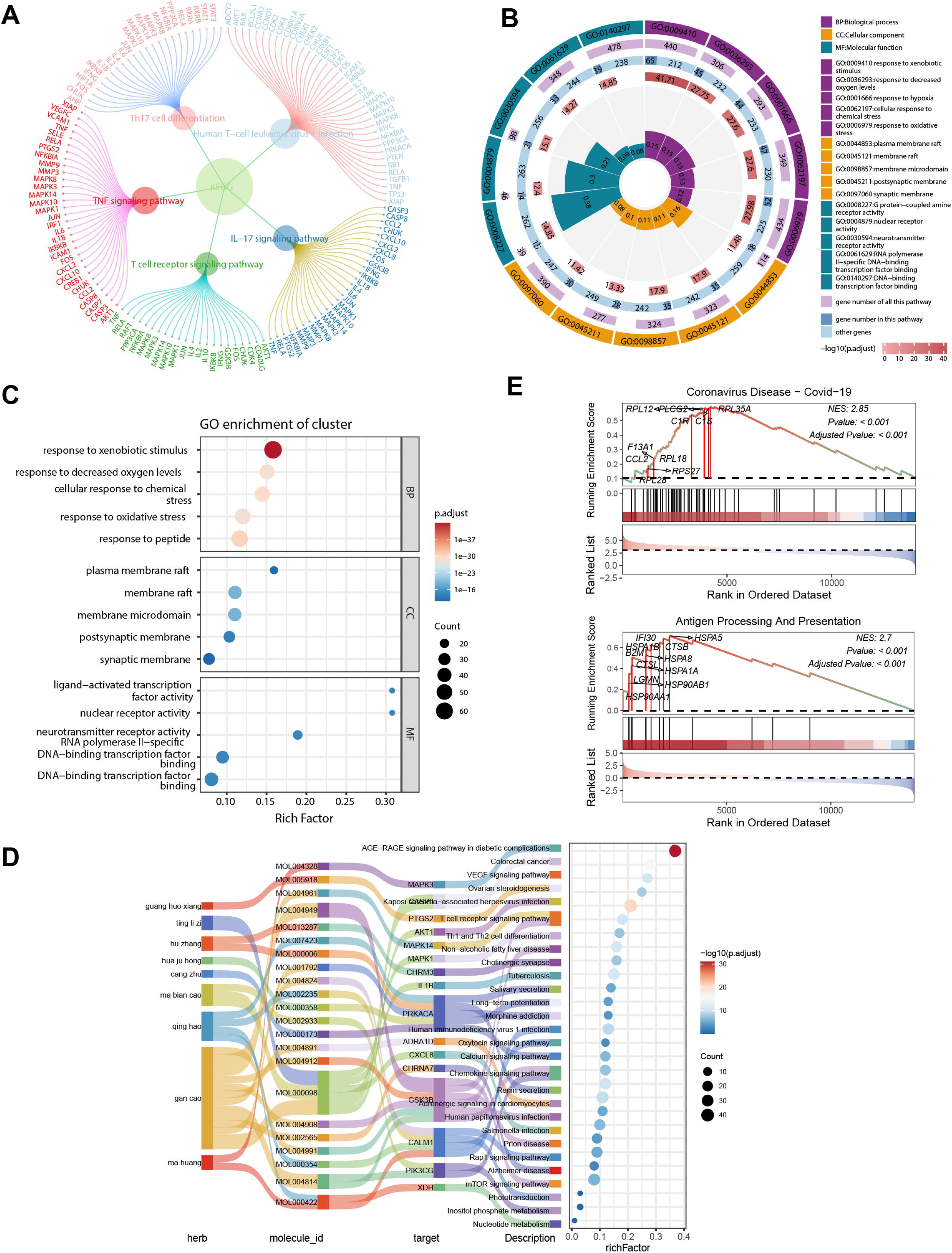
Pathway enrichment analysis and joint analysis of traditional Chinese medicine compound. (A) Circle diagram branches display signaling pathways and genes on each pathway. (B) GO enrichment circle diagram. The first circle represents the pathway id; the second circle represents all the genes in the pathway; the dark color in the third circle represents the enriched genes in the corresponding pathway, and the light color represents the number of other genes involved in this enrichment. The depth of the color in the fourth circle corresponds to -log10 (adjusted *P*-value); the fifth circle (polar coordinate histogram) display GO classification with rich factor. (C) Bar chart showing GO classification results. (D) Sankey and dot graphs showed the results of herb-ingredient-target-pathway network. (E) GSEA enrichment analysis of lung tissue sequencing results of COVID-19 patients.

Additionally, the GO data could be based on biological processes (BP), cellular components (CC), and molecular functions (MF) for categorized displays. Mainly, four functions such as GO bar plot, GO dot plot, GO lollipop plot, and GO circle plot are utilized to present information via bar graph, bubble graph, lollipop graph, and chord graph (Figures 4B, 4C) respectively. We have color-coded and layout-adjusted these functions so they can be displayed using data from KEGG, GO, and other data frame types. Compounds contain drugs that influence pathways by acting on targets via components. To distinctly elucidate the relationship between drugs and pathways, we introduced the tcm_sankey_dot function and tcm_alluvial_dot function. These illustrate the connection between herbs-ingredients-targets-pathways using a combination of Sankey diagram and bubble chart (Figures 4D).

GSEA analysis uses the latest version of human KEGG data sets(347 signaling pathways) and mouse KEGG data sets(348 signaling pathways), hallmark gene sets(human/mouse: 50/50 signaling pathways), curated gene sets (human/mouse: 7233/2664 signaling pathways) and GO gene sets(human/mouse: 16008/10754 signaling pathways)[14]. GSEA analysis can display a single pathway or multiple pathways and can also mark significantly changed genes (Figure 4E). The GSEA data set comes from GSE171524 [15].

### Protein-protein interaction analysis

To determine the interactions among the proteins in the network, we employ PPI analysis. Here, the targets of drug components are extracted and input into the STRING database[16] before downloading the respective data. The PPI net plot implements automatic color adjustment and various layout algorithms. For example: Kamada-Kawai layout algorithm, circle layout algorithm, multi-dimensional scaling, etc. Display graphics include but are not limited to network diagrams, circle diagrams, scatter diagrams, tree diagrams, etc. (Figure 5A).

**Figure. 5.**
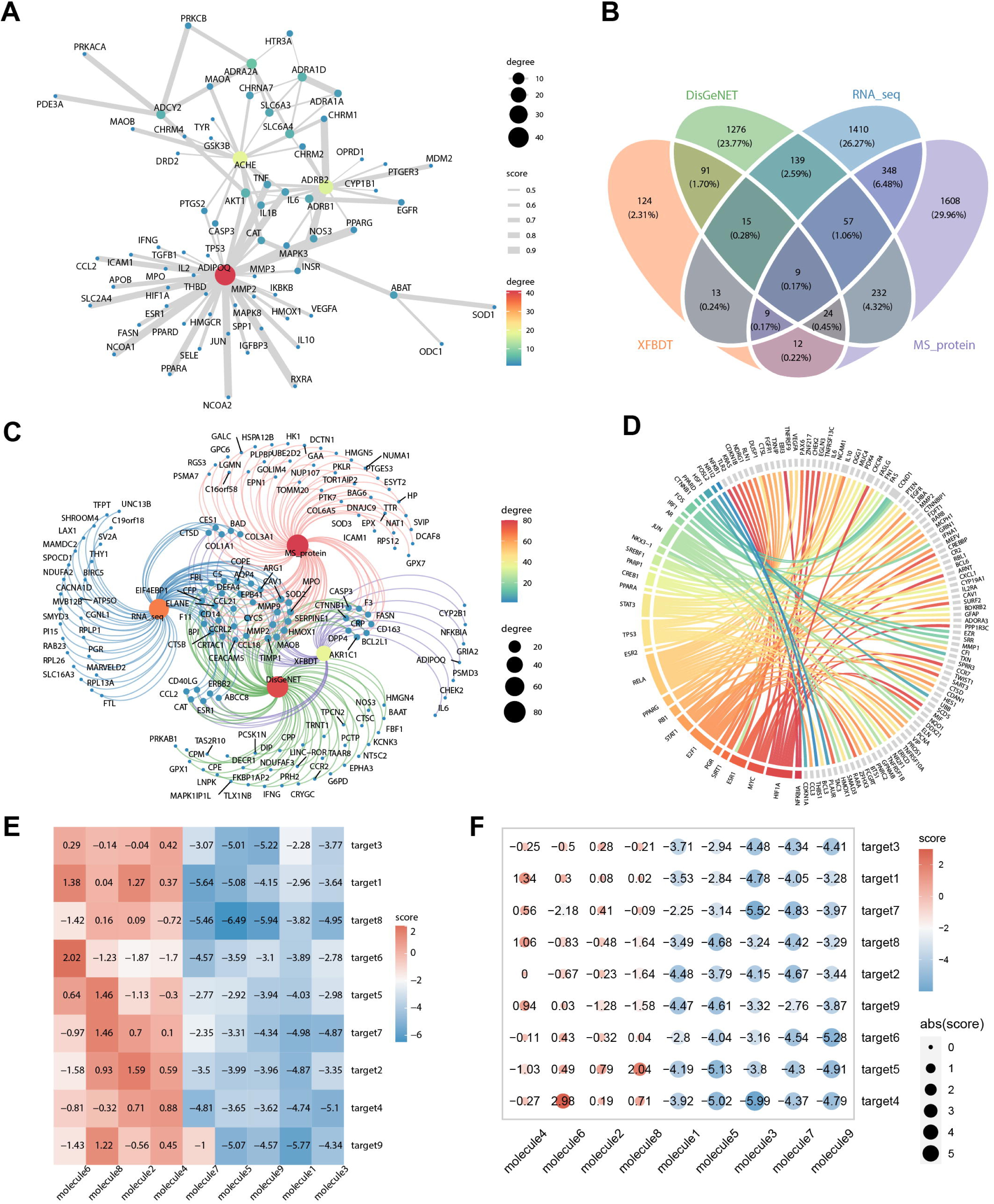
Other functions. (A) Protein-protein interaction (PPI) network. The degree of a node was defined as the number of its connected edges. Score indicates the protein interaction confidence score. (B) Venn diagram showing the intersection of XFBD with disease database and omics sequencing results. (C) Venn network diagram presentation displays elements and the collections they belong to in an interactive network diagram. (D) Transcription factors and the target genes they regulate. The transcription factor is on the left side, and the target gene is on the right side. (E-F) Heat map of simulated molecular docking results. Score indicates the score of the docking result.

### Interaction analysis of drug targets and disease targets

To illustrate the correlation between drugs and diseases, we developed the Venn plot and Venn net plot, which can portray the data in the form of a Venn diagram (Figure 5B). The Venn plot displays shared targets from multiple data sources and thus helps to refine the screening process, thereby aiding in identifying the actual drug. In the Venn net plot, the Kamada-Kawai layout algorithm was used, which can automatically adjust the layout of the graph. Our updated Venn net plot not only shows the content of each intersection, but also allows the user to selectively display intersection content based on the degree of intersection (Figure 5C). In addition, the Venn net plot can also draw network diagrams. The results of the Venn diagram can be obtained through the venn_results button.

### Transcription factor identification

TFs control chromatin and transcription by recognizing specific DNA sequences and forming a complex system that mediates genome expression. TFs are key molecules in the regulation of gene expression. We have also added two more functions to the TCMNP database: TF filter and TF circle plot (Figure 5D). The TF filter function can automatically identify transcription factors and their target genes in human or mouse genes, while the TF circle plot presents these interactions in a visually intuitive circle plot.

### Visualization of molecular and protein docking scores

The interaction between drug ingredients and proteins can be preliminarily predicted through molecular docking. The dock plot has been constructed to present the molecular docking scoring results, which enables preliminary prediction of the interaction between drug ingredients and proteins. The dock plot provides two display formats-heat map and bubble map and facilitates cluster analysis to assist in screening the drug-molecule pairs (Figure 5E).

## Discussion

Network pharmacology is a powerful tool for elucidating the mechanisms behind TCM treatments for various diseases. The TCMNP database provides a series of functions for managing and visualizing network pharmacology data. Compared to other visualization tools, TCMNP simplifies the data analysis process and realizes a complete set of process operations from Chinese herbal compound composition, component and target automatic screening, enrichment analysis visualization, protein interaction and TF and target gene screening. TCMNP provides a more targeted interactive visualization of network pharmacology data. Graphs in TCMNP facilitate the use of ggplot2 for visualization, making it easier for the community to apply further modifications. Moreover, TCMNP harmonizes data structures and can be incorporated into divergent pipelines, including transcriptome and proteome sequencing, for the analysis of drug-treated data, beyond its traditional application in network pharmacology data analysis.

In the process of TCM research, many network pharmacology analysis platforms have emerged, which has promoted the role of network pharmacology in TCM research. At present, there are certain limitations in the process of TCM network pharmacology analysis. First, the components contained in TCM have not been fully determined, which leads to bias and incomplete inferences. Secondly, the effect of dose on target protein is not taken into account. Then, the drug target database targets only reveal the interaction with proteins and lack the connection with RNA and DNA. Finally, due to physical and chemical reactions during the decoction process, TCM compound will generate new complexes or new substances, which are not available for analysis by current network pharmacology. Because of the complexity of TCM compound, exploring the mechanism of TCM compound treatment on disease is still a major challenge. Specifically, establishing a reliable data source, accurate algorithm simulation of drug molecule binding, and scientific verification are necessary conditions to effectively overcome these limitations. By addressing these challenges, researchers can improve the accuracy and reliability of TCM network pharmacology analysis, thereby more comprehensively exploring the effect of TCM in treating diseases and promoting the modernization of TCM.

TCMNP database is freely available and benefits from ongoing maintenance and updates (https://tcmlab.top/tcmr/). the source code necessary to reproduce all plots available is hosted at GitHub (https://github.com/tcmlab/TCMNP). This information also includes details on the datasets utilized in the examples.

## Conclusion

In this work, we developed the TCMNP database and TCMNP package for network pharmacology data analysis and visualization. We believe TCMNP can aid in the comprehensive analysis of the pharmacological workings of TCM in disease treatment and improving our understanding of Chinese herbal compounds and disease.

## Abbreviations

BP: biological processes
CC: cellular components
COVID-19: Corona Virus Disease 2019
ETCM: Encyclopedia of TCM
GO: Gene Ontology
GSEA: Gene Set Enrichment Analysis
KEGG: Kyoto Encyclopedia of Genes and Genomes
MF: molecular functions
PPI: protein-protein interaction
TCM: traditional Chinese medicine
TCMNP: traditional Chinese medicine network pharmacology
TCMSP: TCM Systems Pharmacology Database
TF: transcription factor
TNF: tumor necrosis factor
TRRUST: Transcriptional Regulatory Relationships Unraveled by Sentence-based Text mining
XFBD: xuan fei bai du fang

## Ethics approval and consent to participate

Not applicable

## Consent for publication

All authors agree to publish this article.

## Author Contributions

Jin-kun Liu, Jing Feng, Bin Wu and Min Ying designed the project and supervised the project; Jin-kun Liu and Jing Feng performed all the codes. Jin-kun Liu and Min Ying wrote the manuscript with contributions from all authors; Bin Wu revised the manuscript. All authors read and approved the manuscript.

## Acknowledgments

We are grateful to Prof. Yonghua Wang (Northwest University, Northwest University, China), Prof. Guangchuang Yu (Southern Medical University), Dr. Zuguang Gu (German Cancer Research Center), Dr. Siyuan Hang (Beijing University), Dr. Jianming Zeng (University of Macau), and all the members of bioinformatics team, biotrainee, for generously sharing their experience and codes.

## Availability of data and material

All data and codes generated during the study are available from https://tcmlab.top/tcmr/ and https://github.com/tcmlab/TCMNP.

## Supplementary material

Supplementary table S1. TCMNP database upload file format.

Supplementary code S1. The code used in the article and the main functions of the TCMNP package.

Supplementary video 1. TCMNP package usage guide.

Supplementary video 2. TCMNP database usage guide.

---

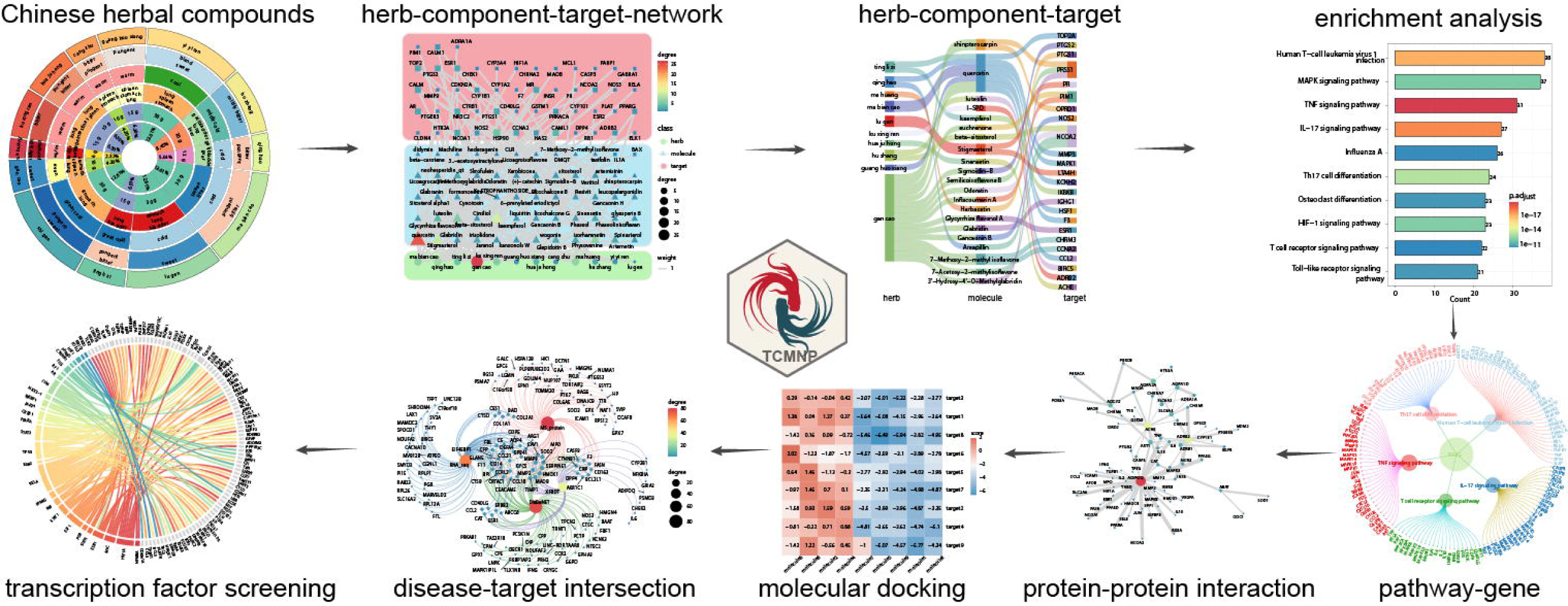

